# Stochasticity of replication fork speed plays a key role in the dynamics of DNA replication

**DOI:** 10.1101/696625

**Authors:** Razie Yousefi, Maga Rowicka

## Abstract

Eukaryotic DNA replication is elaborately orchestrated to duplicate the genome timely and faithfully. Replication initiates at multiple origins from which replication forks emanate and travel bi-directionally. The complex spatio-temporal regulation of DNA replication remains incompletely understood. To study it, computational models of DNA replication have been developed in *S. cerevisiae*. However, in spite of the experimental evidence of replication speed stochasticity, all models assumed that replication fork speed is constant or varies only with genomic coordinates. Here, we present the first model of DNA replication assuming stochastic speed of the replication fork. Utilizing data from both wild-type and hydroxyurea-treated yeast cells, we show that our model is more accurate than models assuming constant fork speed and reconstructs dynamics of DNA replication faithfully starting both from population-wide data and data reflecting fork movement in individual cells. Completion of replication in a timely manner is a challenge due to its stochasticity; we propose an empirically derived modification to replication speed based on the distance to the approaching fork, which promotes timely completion of replication. In summary, our work discovers a key role that stochasticity of the fork speed plays in the dynamics of DNA replication. We show that without including stochasticity of fork speed it is not possible to accurately reconstruct movement of individual replication forks, measured by DNA combing.

**Author summary:** DNA replication in eukaryotes starts from multiple sites termed replication origins. Replication timing at individual sites is stochastic, but reproducible population-wide. Complex and not yet completely understood mechanisms ensure that genome is replicated exactly once and that replication is finished in time. This complex spatio-temporal organization of DNA replication makes computational modeling a useful tool to study replication mechanisms. For simplicity, all previous models assumed constant replication fork speed. Here, we show that such models are incapable of accurately reconstructing distances travelled by individual replication forks. Therefore, we propose a model with a stochastic replication fork speed. We show that such model reproduces faithfully distances travelled by individual replication forks. Moreover, our model is simpler than previous model and thus avoids over-learning (fitting noise). We also discover how replication speed may be attuned to timely complete replication. We propose that fork speed exponentially increases with diminishing distance to the approaching fork, which we show promotes timely completion of replication. Such speed up can be e.g. explained by a synergy effect of chromatin unwinding by both forks. Our model can be used to simulate phenomena beyond replication, e.g. DNA double-strand breaks resulting from broken replication forks.

## Introduction

DNA replication in eukaryotic cells is highly regulated to ensure that the whole genome is duplicated correctly and completely before cell division [1]. Replication initiates at specific sites, termed origins of replication. Origins are prepared to be activated (i.e. fired) with the assembly of a pre-replication complex, through a process termed licensing, during the G1 phase [2]. Replication origins are licensed in excess and during the subsequent S phase a subset of origins initiate replication. Two forks emanate and elongate bi-directionally from each active origin, the rest of the licensed origins are passively replicated by the forks emerging from the neighbor origins [3, 4]. In the budding yeast *Saccharomyces cerevisiae*, DNA replication initiates from *∼*400 origins with known genomic coordinates [5]. Origin activation is stochastic in individual cells [6, 7], but chronological order of origin activation is reproducible population-wide. This flexibility in origin activation is essential in response to DNA damage and adaption of replication to gene expression [8, 9]. Upon origin activation, replication forks are formed and progress along the chromosome until they meet another fork moving in the opposite direction. High-throughput experimental data, which have been used to study the dynamics of DNA replication, allow the measurement of average replication time and average fork speed, but mask the cell to cell variations in these parameters [10]. Distances travelled by individual replication forks *in vivo* can be visualized and measured using DNA combing. However, DNA combing does not provide the genomic coordinates, and complexity of spatio-temporal regulation of replication makes interpretation of these data difficult. As a result, computational models are necessary to analyze the mechanism of DNA replication and understand how regulation of origin activation and fork elongation impact its dynamics.

Substantial stochasticity of replication fork speed has been observed in *in vitro* biophysical studies of individual forks [11] and in DNA combing and 2D gel analysis in S. cerevisiae [11–16]. Nevertheless, previous models assumed either that fork speed was constant globally [7, 17–29] or constant in specific genomic regions [30, 31]. Moreover, previous models used only population-wide data and typically employed origin-to-origin comparison used for validation and parameter selection [22–24, 31]. Such an approach can elucidate information about origin average firing time and efficiency (i.e. percentage of cells in which origin is fired), but it cannot distinguish between variability in the fork speed and the stochasticity of origin firing time.

Here, we present Repli-Sim, a probabilistic numerical model for DNA replication, which simulates DNA replication in *S. cerevisiae* genome-wide assuming stochastic replication fork speed. Repli-Sim includes local parameters specific to each origin inferred from experimental data and global parameters assigned to origins using a Monte Carlo method, which are optimized through a genetic algorithm. To validate our model we used DNA combing data showing distances traveled by individual replication forks (DNA tracks) and genome-wide DNA copy number data. We show that stochasticity in the fork speed is key to reconstructing dynamics of DNA replication in single cells measured by DNA combing. We also show that constant speed models, such as previously used, are incapable of accurately reconstructing distribution of distances traveled by individual replication forks (DNA tracks). We also report the observation, based on three independent datasets, that an individual fork speed depends on the distance to the approaching fork. We show that such modification of the fork speed promotes timely completion of the replication, which is a challenge due to stochastic nature of replication.

## RESULTS

We will use both a single origin of replication and the whole genome to show how the variance of fork progression rate impacts DNA track distribution. For single origin of replication analysis, we will illustrate a significant increase in the difference between variable and constant fork speed at later times during S phase, representing a more dominant effect of fork progression rate variability at longer times, while the average length of the DNA track remains comparable in all models. For genome wide analyses, Repli-Sim is utilized to derive the DNA tracks for both normal and HU-induced cells and it is shown how taking into account variability in replication fork speed impacts the dynamics of replication and distribution of DNA tracks.

### Simulations of DNA replication (Repli-Sim)

Repli-Sim, is a probabilistic numerical model designed to study the dynamics of DNA replication. During S-phase, origins of replication are activated and DNA tracks (continuous distances covered by replication forks, Fig. 1) are formed and elongated throughout the genome until the whole DNA is replicated. In Repli-Sim, coordinates *x* of replication origins are derived from experimental data and filtered using a database of replication origins, OriDB [5]. As shown in Fig. 1, two forks are formed and elongate bidirectionally across the genome to form DNA tracks (Δ*x*). For each origin *i* in a cell population, at time *t*_*exp*_ (measured from the beginning of DNA replication), we derive the distribution of Δ*x* based on two assumptions. First, the firing time of the origin, 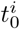, is derived from a normal distribution with a mean firing time 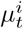(specific to that origin) and with global standard deviation *σ*_*t*_:

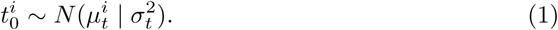

**Fig 1.**
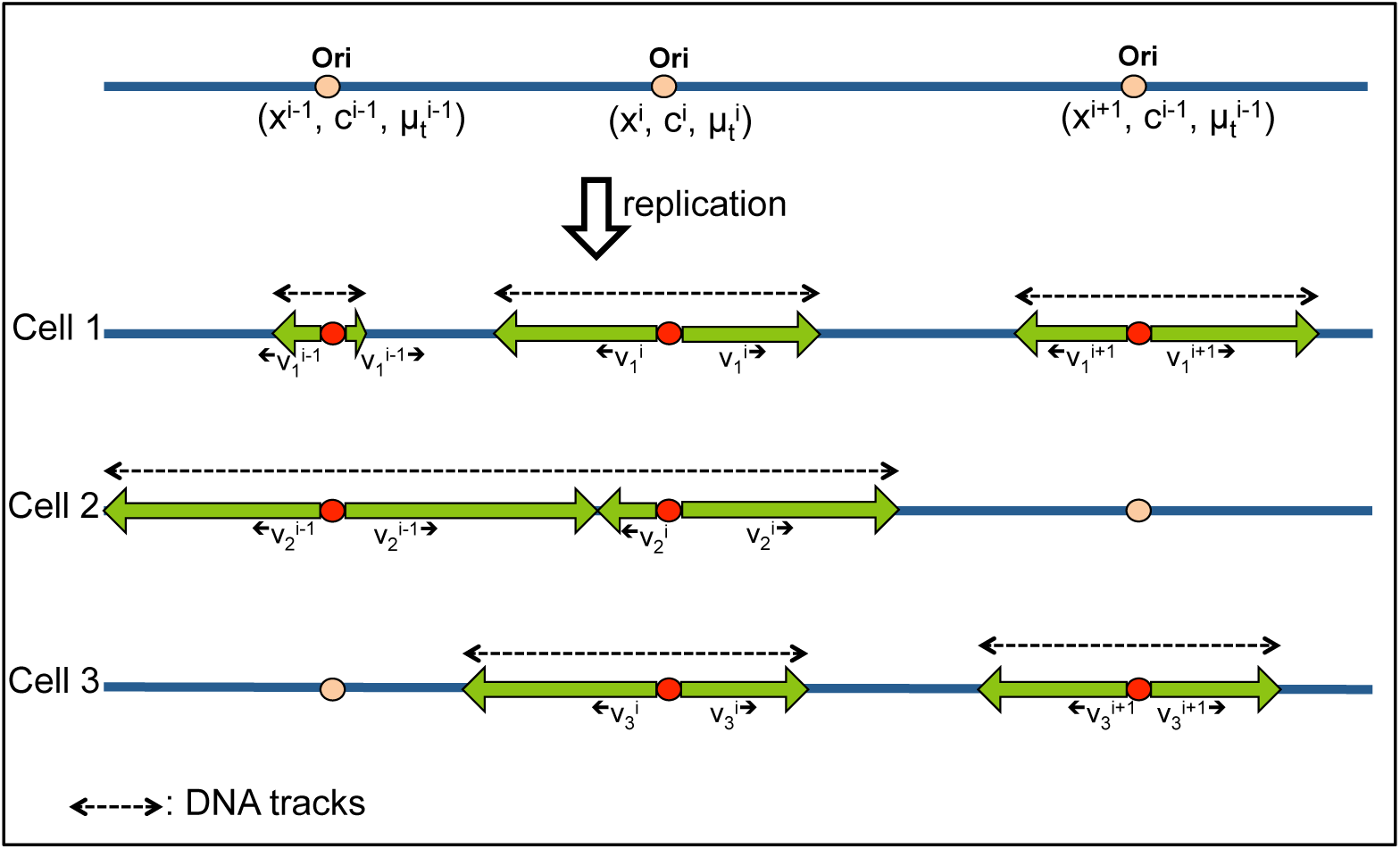
A schematic of the mechanism of DNA replication encoded in Repli-Sim. Repli-Sim includes local origin parameters (position *x*^*i*^, competence *c*^*i*^, and mean firing time 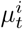) and global parameters (firing time variance *σ*_*t*_, mean fork speed *µ*_*v*_ and its variance *σ*_*v*_). When an origin of replication activates, two forks are formed and elongate bidirectionally until they meet an approaching fork. The continuous length of the replicated DNA (Δ*x*, DNA tracks) are shown with the dashed lines.

Mean origin firing time 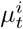 is derived from experimental data, as described in Methods. Second, individual forks are assigned with different speeds, *v*^*i*^, derived from the same probability distribution with a mean speed, *µ*_*v*_ and standard deviation *σ*_*v*_:

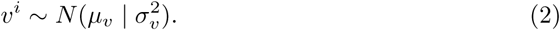

A probability of origin licensing *c*^*i*^ (a priori probability of origin activation) is assigned to each individual origin as a random number between the experimentally measured frequency of that origin activation and 1. Then, a Monte-Carlo method is used to generate activation time 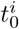 for an origin *i* from a Gaussian probability distribution with an experimentally estimated mean activation time 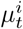 specific to that origin (Methods), and a global standard deviation *σ*_*t*_, same for each origin. Individual forks progress with different speeds, constant for each fork, generated using a Monte-Carlo method from a Gaussian probability distribution with a global average speed *µ*_*v*_ and standard deviation *σ*_*v*_.

### Impact of fork speed stochasticity on the dynamics of DNA replication

First we illustrate the impact of variance in fork speed on distribution of the DNA tracks by analyzing single origin of replication. In Figure 2 we show the distribution of DNA tracks (Δ*x*) for constant (*σ*_*v*_ = 0) and variable (*σ*_*v*_ ≠ 0) fork speed and for single origin. The difference between variable and constant fork speeds are especially pronounced later in S phase, while the average length of the DNA track remains similar for both models. We have shown (Methods Eq. 6)

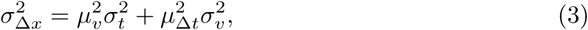

**Fig 2.**
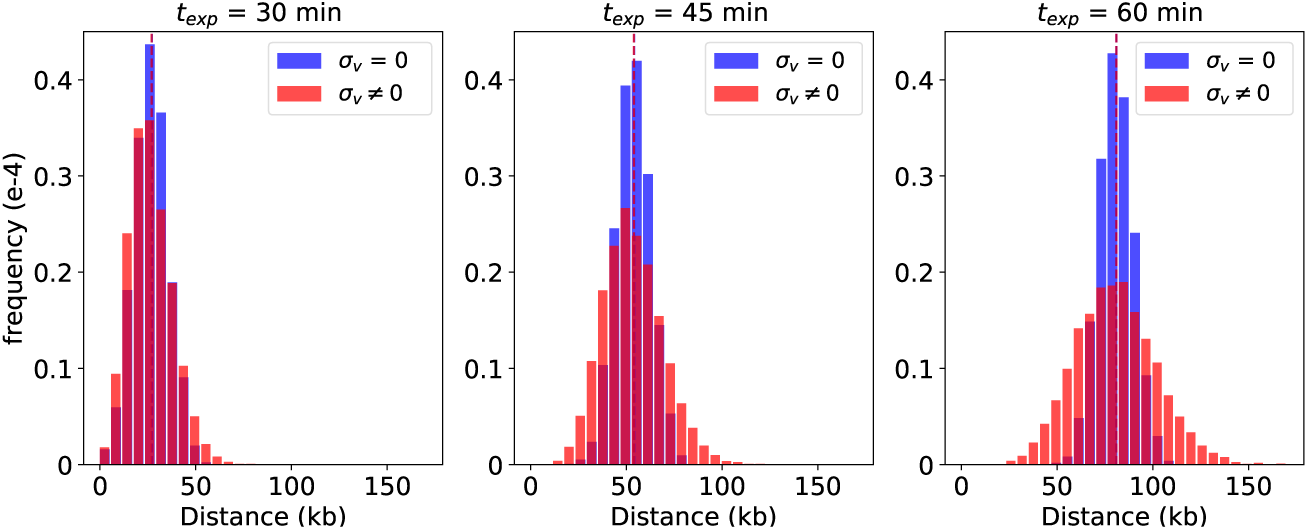
The impact of stochasticity of replication fork speed on the distribution of DNA tracks. The distributions of DNA tracks for constant (*σ*_*v*_ = 0, blue) and variable (*σ*_*v*_ ≠ 0, orange) replication fork speeds at three different time points within the S phase: 30, 45, and 60 minutes. The differences in DNA track distributions between constant and variable speed models become most pronounced at later times.

which implies that stochasticity of distribution of DNA tracks (*σ*_Δ*x*_) not only depends on the average fork speed (*µ*_*v*_) and average firing time (*µ*_Δ*t*_) but also on their degree of randomness (*σ*_*v*_ and *σ*_*t*_). On the other hand, considering Eq. 3, assuming a constant fork speed (*σ*_*v*_ = 0), the second term 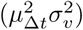 vanishes. Therefore, fitting *σ*_Δ*x*_ using constant speed models will lead to over estimation of *σ*_*t*_*µ*_*v*_. Since the average fork speed is relatively easy to determine experimentally, fitting distribution of DNA tracks using constant speed models will result in artificially increased stochasticity of origin firing, manifesting itself e.g. by known late origins to fire early in S phase, as if they were early origins, as in previous models [23]. We discuss stochasticity of origin firing time in more detail elsewhere (Yousefi et al. in preparation).

### Examining fork speed stochasticity in wt yeast cells

To investigate whether fork speed is stochastic or constant in wild-type (wt) yeast cells, we used time course DNA sequencing data [23]. These experimental data were taken every 5 minutes between minute 15 to 40 during S phase and include mean firing times and efficiencies of the origins, which we utilized in our analysis. Other parameters including *σ*_*t*_, *σ*_*v*_, *µ*_*v*_, and time of observation *t*_*exp*_ were selected by Repli-Sim through identifying the best-fitting model via simulations. For both constant and stochastic fork speed models, the simulations were performed for > 5000 sets of parameters selected by a genetic algorithm. For each parameter set we used least square difference (Fig. 3(a)) to identify parameters leading to the distribution of DNA tracks most closely resembling the distribution of DNA tracks obtained from experimental data. Figure 3(b) presents the results for best fitting parameters for both constant and stochastic fork speed models and shows that a model with stochastic fork speed fits the experimental data best. The best fitting model exhibits considerable relative stochasticity of fork speed (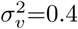 (kb/min)^2^). Strikingly, that same relative stochasticity (*σ*_*v*_*/v*) of replication fork speed that we derived from simulations was observed in in vitro studies of individual replication forks in another organism [11]. The average replicated distance in the stochastic speed model is comparable with that of the experimental data (105 kb). In addition, the replication fork speed and *t*_*exp*_ derived from simulations for the variable fork speed model (1.5 (kb/min), 42 (min)) are more consistent with experimental data (1.6 (kb/min), 42 (min)) than those obtained from the best constant speed model (1.4 (kb/min), 50 (min)).

**Fig 3.**
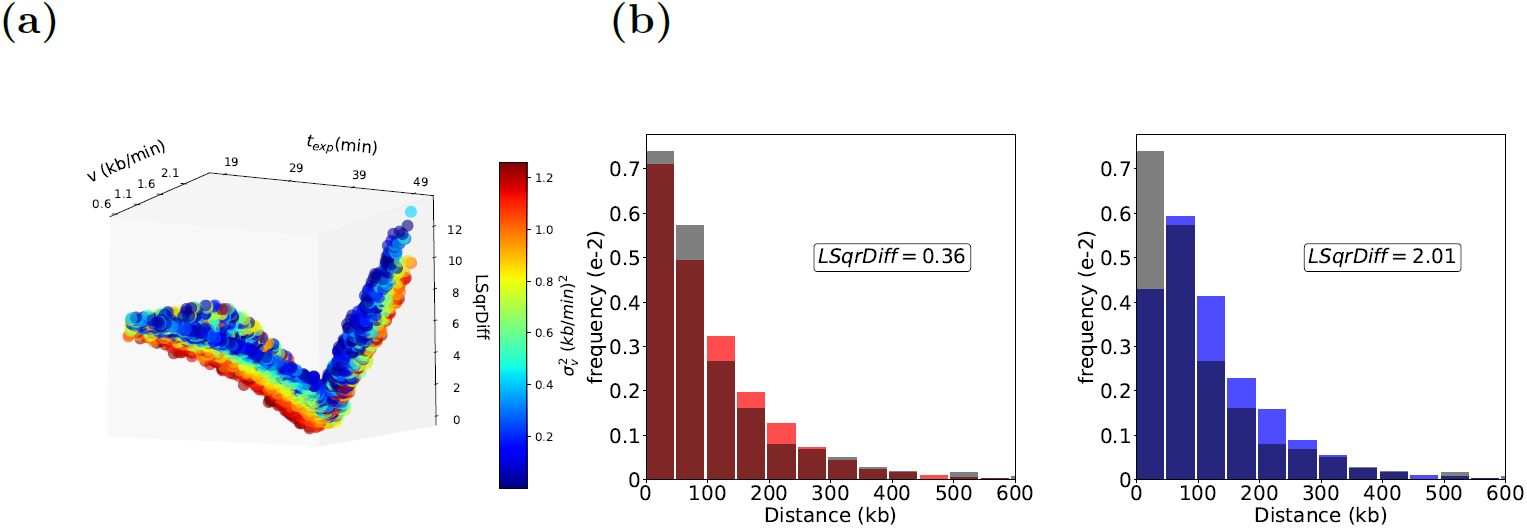
Model selection by Repli-Sim for models with constant and variable fork speeds. (a) All models (parameter sets) considered. The fork progression speed (*v*) and experimental time (*t*_*exp*_) are shown on the horizontal axis, least square difference (LSqrDiff) with experimental data (the lower the better) is shown on the vertical axis. Stochasticity of replication fork speed 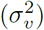 is color coded, as shown in the color-bar. Best models (smallest LSqrDiff value) are more stochastic. The best selected constant speed model had parameters 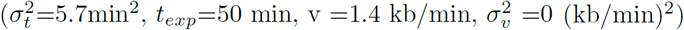 and the best variable speed model was 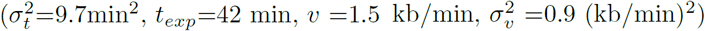. The *t*_*exp*_ and fork speed from experimental data are 40 min and *v* = 1.6 (kb/min), which are more compatible with the variable fork progression model. (b) The distribution of DNA tracks for both best constant (blue) and variable (orange) models are shown along with the distribution of DNA tracks from experimental data (gray), which shows a better fit for a variable speed model. The average distance traveled in the variable speed model is compatible with the experimental data (105 kb).

### Examining fork speed stochasticity in hydroxyurea-treated wt yeast cells

Hydroxyurea (HU) is an inhibitor of the ribonucleotide reductase (RNR), an essential enzyme for catalyzing the production of deoxyribonucleotide triphosphates (dNTPs), the building blocks of DNA. As a result, HU treatment depletes dNTPs thus slowing replication fork progression and making HU-treated cells an interesting case to study. To examine the stochasticity of fork speed and its impacts on replication in HU-treated cells, we used experimental DNA track data from HU-treated wt yeast cells studied in [32]. Mean origin firing time for each individual origin was derived as described in Methods. Similar to the previous analysis, for both constant and stochastic fork speed models, the simulation utilized 5000 sets of parameters selected randomly by a genetic algorithm, and for each parameter set the distributions of DNA tracks were derived and compared with the experimental DNA track distribution by calculating the LSqrDiff between the distributions binned with 1*kb* bin size. We first identified a group of best fitting models (LSqrDiff < 0.65), and then as the final model we selected the model with a total number of active origins consistent with that of the experimental data (280 ± 10). It is important to note that stochasticity of firing time *σ*_*t*_ impacts origin usage. A smaller *σ*_*t*_ is equivalent to more narrow firing time, which leads to activation of fewer late origins early in S phase as compared to a larger *σ*_*t*_, as we discuss elsewhere (Yousefi et al. in preparation). Indeed, the dysregulation of origin activation has been observed in various conditions [32–35], which could be explained by increasing stochasticity of firing time of origins of replication *σ*_*t*_. All the parameter sets from simulations are presented in Fig. 4(a), where models with a smaller LSqrDiff value (i.e. better fitting), exhibit more stochasticity in fork speed (higher *σ*_*v*_). The best models, among models with the lowest LSqrDiff value, are selected, using numbers of active replication origins, as described above. In Fig. 4(b) we compare experimental and best-fitting simulated distributions of DNA tracks for constant speed and stochastic speed models. Same as for untreated wt yeast cells, the stochastic model fits the data much better for HU-treated cells.

**Fig 4.**
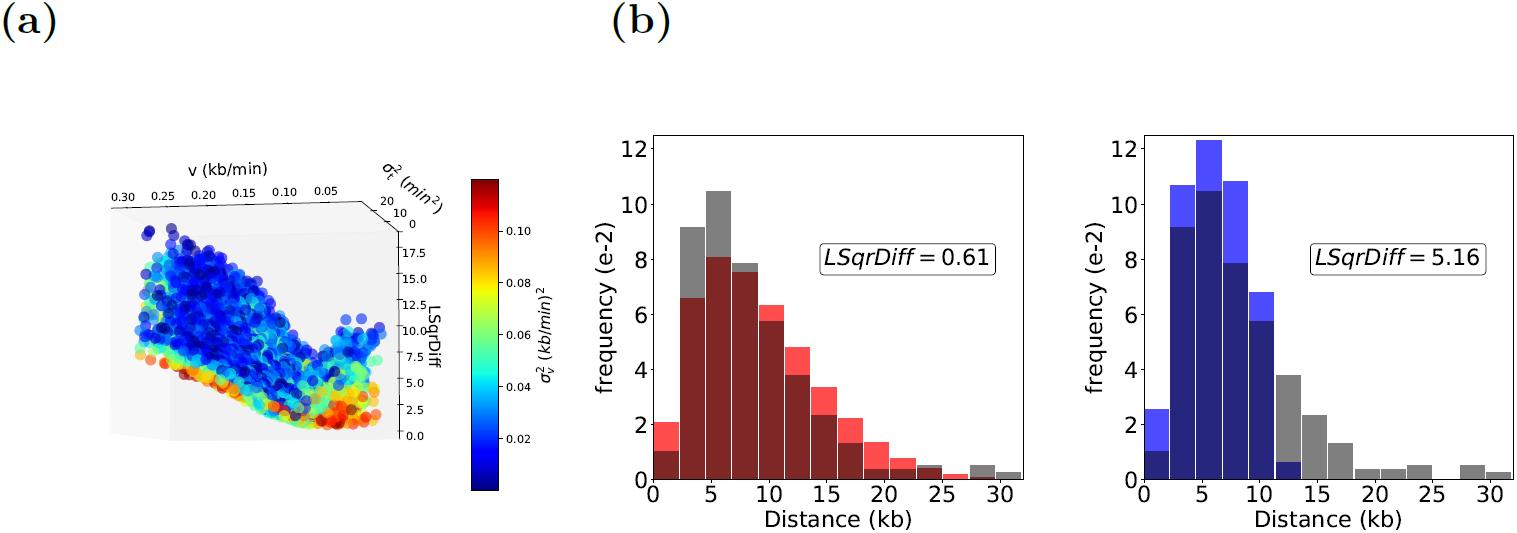
Results of Repli-Sim for HU-treated cells for both constant and variable fork speed models. (a) All models (parameter sets) considered. Fork speed (*v*) and stochasticity in firing time of the origins (*σ*_*t*_) are shown on the horizontal axes and least square difference (LSqrDiff) with the experimental data of [32] is shown on the vertical axis, stochasticity of replication fork speed 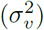 is color-coded (side bar). Best fitting models (smallest LSqrDiff) are characterized by more stochastic fork speed. Best fitting constant speed model is 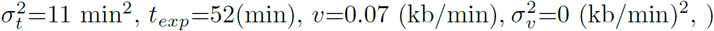 for the best selected variable speed model is 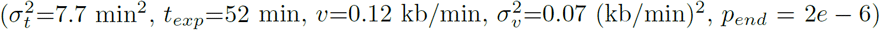. (b) The distribution of DNA tracks for both best constant (blue) and variable speed (orange) models are shown, along with the DNA tracks from experimental data (gray), which shows a better fit for a variable speed model.

### Comparison with the previous work

To compare Repli-Sim fairly with the most current published model of DNA replication [23], we obtained new DNA track data to avoid using data on which our model was trained previously. In the Hawkins et al. model, unlike in our model, origins have not only individual assigned firing time, 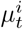,but unlike in our model, each origin has its individual firing speed stochasticity, 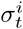, resulting in 814 model parameters. To highlight the impact of stochasticity of replication speed on the accuracy of reconstruction of DNA replication dynamics, we prepared a simplified Repli-Sim model, where origin firing time is stochastic (not empirically derived as previously). Such a simplified Repli-Sim model only has 5 parameters (mean and variance of origin firing time, mean and variance of fork speed, time of taking measurement), in addition to origin coordinates, considered known. As Figures 5 shows, even such a simplified model fits the DNA tracks data much better than the more complex Hawkins et al. model [23]. This results highlights the key role stochasticity of replication speed plays in accurately reconstructing dynamics of DNA replication and thus DNA tracks. It is also unusual that the model with fewer parameters fits the data better, which again stresses the importance of including stochastic replication speed in a model, without it, accurate fitting of the data it is not possible.

**Fig 5.**
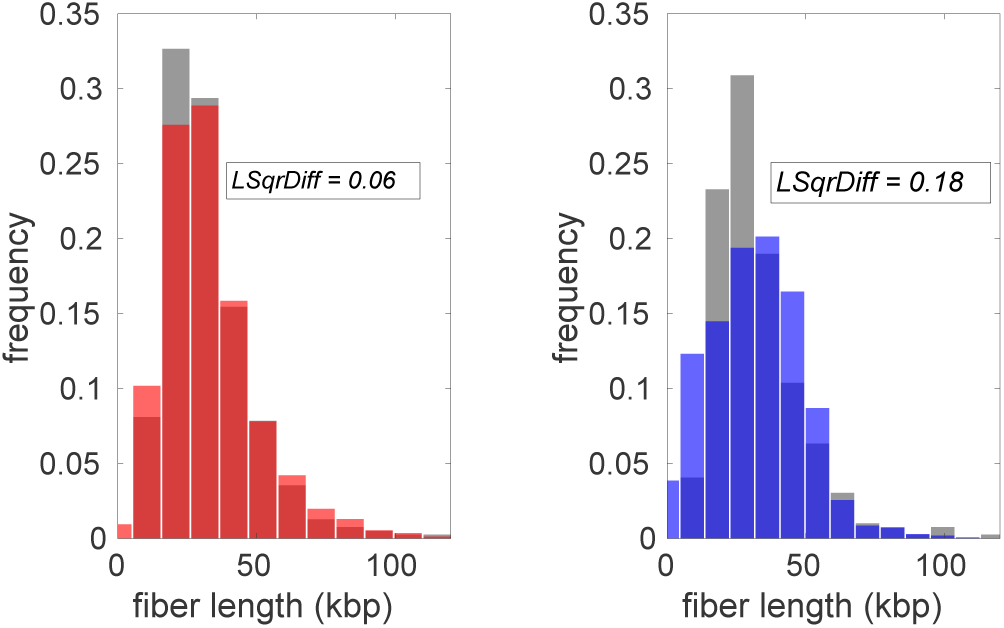
Comparison with the previous model. Our simplified (random origin activation times, but stochastic fork speed) model (orange), reproduces distribution of distances travelled by replication forks measured by DNA combing (gray) much better than previous model (blue).

### Context-dependent variability of fork speed and its impact on the completion of replication

DNA replication dynamics is impacted not only by origin activation, but also by replication fork speed. It has been shown that different cellular conditions (e.g. dNTP levels) impact speed of the fork [32, 36, 37]. Here, we analyzed DNA copy number in three independent data sets ([23], [13] and [12]) and found that fork speed increases dramatically if there is an approaching fork nearby (Figure 6(a)). High correlation between fork speed and the distance to the approaching fork is shown in Figures 6(b-d). This plasticity of fork speed could be the reason for the higher stochasticity of DNA track distribution observed for later firing origins [24]. The mechanistic explanation for this increase in fork speed may be a synergistic effect of DNA unfolding in front of the replication fork, resulting in faster fork progression.

**Fig 6.**
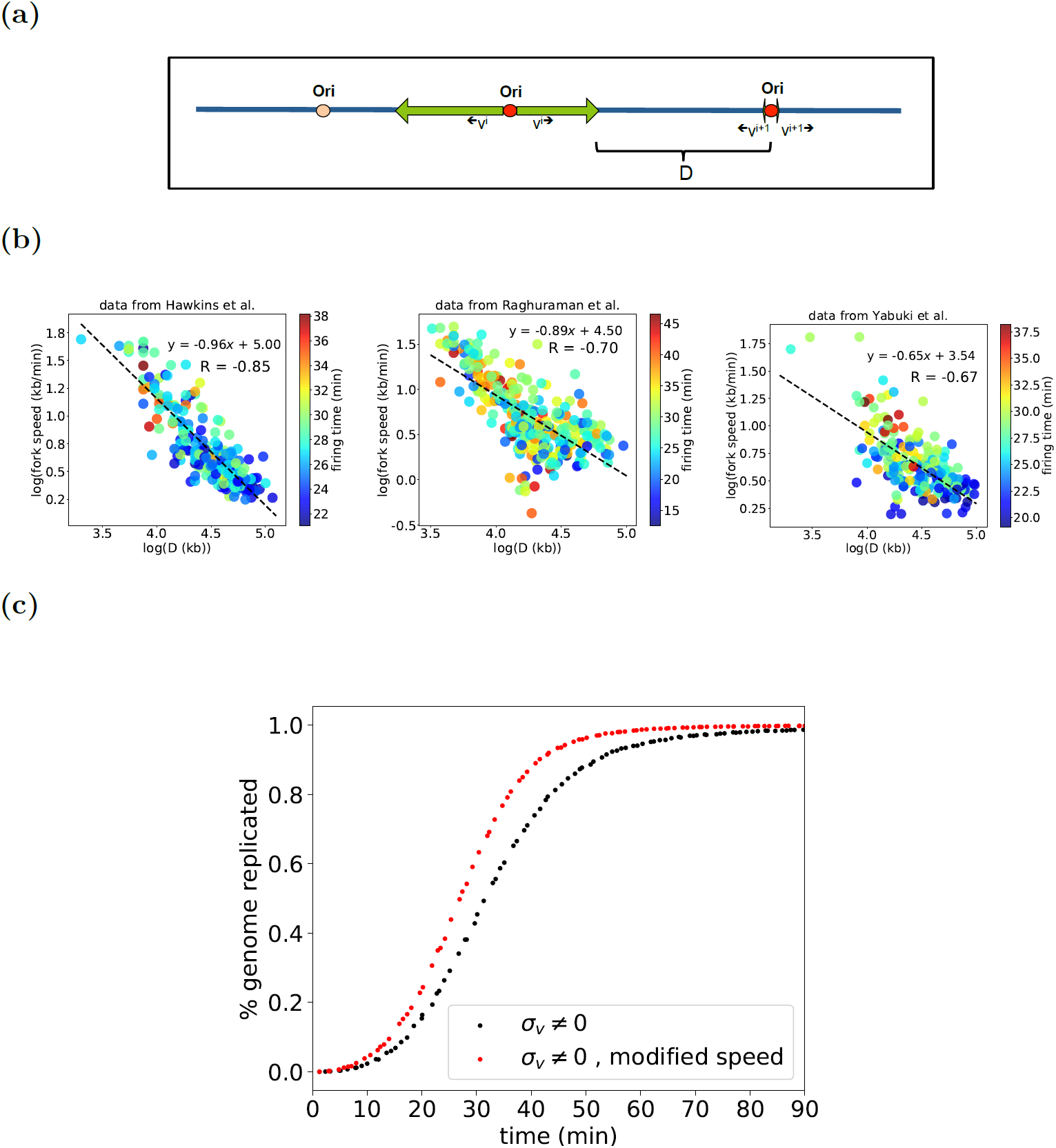
Replication fork speed adjustment based on the distance to the approaching fork. (a) Schematic representation of *D*, the distance from an emerging fork to an approaching fork. (b) The strong correlation between *D* and emerging fork speed is observed in three independent data sets [12, 13, 23]. (c) The impact of fork speed modifications on the dynamics and completion of DNA replication. The percent of replicated genome is presented as a function of time for stochastic fork speed models with regular (blue) speed and modified speed based on the distance to the approaching fork (green). The modified fork speed model promotes timely completion of replication.

To examine the impact of the observed increase in fork speed, the fork progression rate in Repli-Sim is modified, based on the our observations, by assigning a fork speed to origins of replication that depends on the distance of that origin to the approaching fork. The stochastic nature of replication leads to the well known random gap problem [38–42], which challenges the completion of DNA replication in a timely manner. All the proposed solutions to address this problem, have been focusing on the regulation of origin activation [39, 40], while regulation of replication fork progression, which impacts the dynamics of replication as well, has not been taken into account. As shown in Fig. 6(d), the random gap problem can be solved by the proposed modification in the fork speed, since such a modified fork speed model promotes the completion of replication, so that in the modified speed model more that 99% of the genome is replicated, while in normal case it takes more than 90 minutes.

In order to verify our assumption, the experimental data for replication timing profile (the time at which 50% of the DNA at specific coordinates is replicated) of chromosome I [23] is compared with replication timing profile of both modified and non-modified stochastic fork speed models. As shown in Fig. 7(a), the modified fork speed model fit the experimental data the best. In addition, replication timing profile for non-modified high speed fork progression model is presented, which shows that deriving the fork speed from replication timing profile is not accurate and can lead to over estimation of fork speed progression as in [12]. DNA copy number profiles for three different time points (20, 30, and 40 minutes) are presented in Fig. 7(b), which shows the differences between the replicated DNA content at different times for both modified and non-modified models.

**Fig 7.**
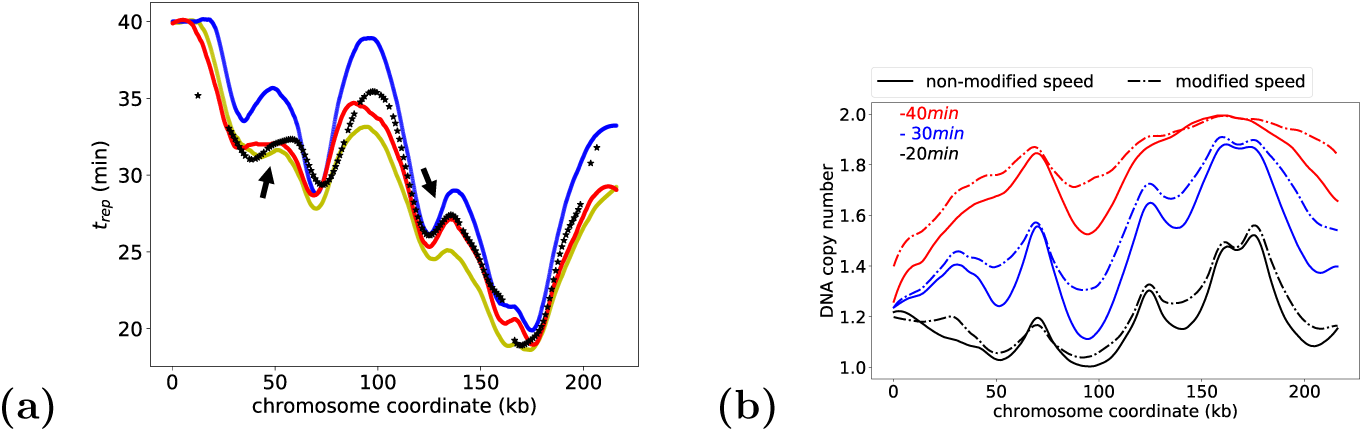
Replication timing profile and DNA copy number of chromosome *I*. (a) Schematic representation of replication timing profile for experimental data ([23]) (black), along with modified (red) and non-modified (blue) fork speed models, which shows a better fit of the experimental data with modified fork speed model. As shown, modified fork speed fits better not only overall replication timing profile, but also in the regions with higher fork speed (smaller slope), indicated by arrows. In order to check if other factors including a high fork speed model can lead to the same observation, the replication timing profile for a much higher (2.3*kb/min*) fork speed is presented (yellow), however it is not compatible with a measured fork speed from time course data (1.6*kb/min* in [23]). (b) DNA copy number at different time points for both modified and non-modified speed models.

**Fig 8.**
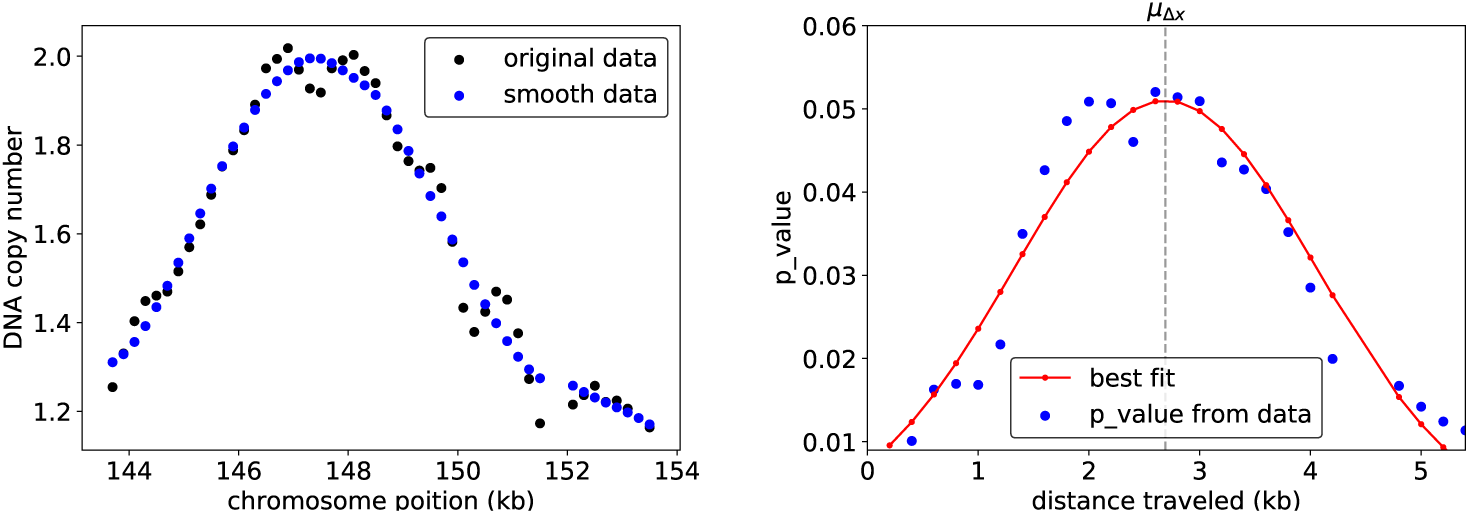
An illustration of the derivation of *µ*_Δ*x*_ for each individual origin. (a) As an example we use the DNA copy number of an early origin located at 147 kb from the beginning of the chromosome I. The distribution of DNA tracks measured from BrdU data [32] is normalized based on the BrdU micro-array DNA copy number of origin ARS305, which was verified by quantitative PCR in the same experimental condition. For smoothing the data, the Savitzky-Golary filter (working through the convolution process) is utilized, because it minimally distorts the original data. Maximum of the smoothed peak indicates the origin position. (b) The data are transformed into probability distribution function (PDF) of Δ*x* and fitted with a Gaussian distribution which peak is assumed to be *µ*_Δ*x*._

## DISCUSSION

### Key role of stochasticity of replication fork speed

Even though the experimental data, both from single-cell biophysical studies of replication fork and from visualizing DNA track in vivo (DNA combing) indicate that replication fork speed is highly stochastic, all published DNA replication models assumed constant replication fork speed or speed depending only on genomic coordinates. Here, we present Repli-Sim, the first model of DNA replication including stochastic replication fork speed. We have shown that Repli-Sim matches DNA tracks travelled by individual forks much better than models with constant speed. To illustrate how important stochastic speed is for accurate DNA track matching, we simplified our model to only 5 parameters and nevertheless obtained better fit with DNA track data than the much more complex Hawkins et al. [23] model, utilizing 814 parameters.

We have shown that standard deviation of the length of DNA tracks (i.e. distances travelled by individual forks), *σ*_Δ*x*_, for each origin can be approximated by the formula:

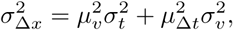

where *µ*_Δ*t*_ is the average time elapsed since the origin was activated. This formula shows why the previous modeling attempts, assuming a constant replication fork speed (*σ*_*v*_ = 0), were not successful: *σ*_*v*_ is an important contributor to *σ*_Δ*x*_, so assuming empirically incorrect *σ*_*v*_ = 0 forces compensation with *σ*_*t*_, resulting in incorrectly derived timing of the origins. Alternatively, if activation time and its standard deviation *σ*_*t*_ is derived origin by origin from the experimental data, and *σ*_*v*_ is assumed zero, it leads to substantially distorted DNA tracks distribution *σ*_Δ*x*_. On the other hand, *µ*_Δ*x*_ is not affected by stochasticity of speed, which is why the stochasticity of speed is most apparent in single-cell derived data such as DNA tracks, where *σ*_Δ*x*_ becomes visible. It is also important to note that a larger stochasticity of firing time *σ*_*t*_ is equivalent to observation of activation of many origins including late origins during early S phase [43].

### Model selection

During model development, a better fit can normally be achieved with an increased number of parameters, which may lead to overly complicated models and over-fitting (learning noise). In information theory, criteria have been developed to help find balance between goodness-of-fit and using parameters sparingly, e.g. Bayesian Information Criterion (BIC): BIC = k log n − 2 log L, where *n* is the number of data points, *k* is the number of parameters, *L* is likelihood and log is natural logarithm. The model with the smallest BIC is considered optimal. BIC penalizes increasing number of parameters if they do not increase goodness of fit sufficiently. Interestingly, when we compared even a simplified version of our model (random origin firing time) with the best previous model, it turned out that our model has both much fewer parameters and fits DNA track data much better (Fig. 5), resulting in dramatically better BIC: 48 for our simplified model and 7310 for Hawkins et al. model (lower values indicate better model). This result highlights the critical role of stochasticity of replication speed (*σ*_*v*_ ≠ 0) in correct modeling. To develop a model with good BIC (i.e. to avoid over-fitting), we assume the same *σ*_*t*_ for all origins and do not attempt to match the data to individual origins. Instead, we transform experimental data into distribution of DNA tracks (estimated distances travelled by forks) and optimize its fit with DNA tracks data generated from simulated conditions. Moreover, using single *σ*_*t*_ instead of > 400 individual *σ*_*t*_(*i*) as in [23], gives clearer insights into changes in replication program in HU-treated cells.

### Challenge of completing replication on time

Our results presented hare and previous experimental results clearly show that replication fork speed is stochastic. Stochasticity of replication speed makes it a challenge to complete replication timely, for constant speed replication will always be completed in finite time. Here, based on observations in three independent data sets, we proposed a potential new mechanism that may ensure timely completion of replication. We hypothesize that the exponential increase in fork speed observed when the approaching fork is very close may be resulting from a synergistic effect of DNA unfolding in front of the replication fork, resulting in faster fork progression. More research is needed to test this intriguing hypothesis.

### Applications and future directions

DNA replication is a complex process, with elaborate spatio-temporal regulation, which remains incompletely understood. Due to this complex regulation of replication, it is difficult to infer the role individual proteins play from genomic copy number variation or DNA track data, since changes in origin activation and replication fork speed can be difficult to distinguish in such data. Here, we present Repli-Sim, a probabilistic model of DNA replication including stochastic replication speed. We have shown that Repli-Sim accurately reproduces experimental data. Moreover, Repli-Sim allows the user to classify experiments in terms of fundamental parameters of replication, such as replication fork speed, its stochasticity, and stochasticity of origin firing. Such presentation allows us to better understand the impact that individual treatments and proteins have on DNA replication, as well as compare conditions in this space of fundamental replication parameters (Yousefi et al. in preparation). Another application of our simulations can be studying replication stress and DNA double-strand breaks (DSBs) originating from broken replication forks. Currently, the numbers of DSBs per cell can be precisely measured genome-wide using qDSB-Seq [44]. However, since replication stress typically substantially changes the replication program, increased numbers of breaks per cell do not have to mean that forks break more often. Therefore, combination of DNA replication simulation by Repli-Sim with the landscapes of DSBs measured by qDSB-Seq, allows deeper insight into how stalled replication forks break and form DSBs as a result of replication stress (Zhu, Biernacka et al., in preparation).

In summary, Repli-Sim is designed to be general and usable with different input data types, in contrast with [23], which is designed to use microarray data only. Here, we showed how Repli-Sim can be used with DNA combing data as an input. Repli-Sim can also use DNA copy number data from sequencing or microarrays as an input, after pre-processing the data to derive DNA track distribution, assuming it is Gaussian. Such pre-processing has an additional advantage that it acts as a smoothing procedure and reduces the noise. Last but not least, Repli-Sim is very fast, simulations of DNA replication in a given condition require testing of 10,000 sets of parameters, which takes only 7 hours to perform on a 16-core, 32-thread 3.1GHz workstation. Therefore, Repli-Sim can be used to infer spatio-temporal organization of replication in variety of conditions, as long as appropriate data is available. Once Repli-Sim derives parameters of a given state, also spatio-temporal organization of replication and later and earlier time-points can be reconstructed. Therefore, Repli-Sim can play a role similar to the role of high-throughput screening in drug discovery: allowing very fast testing of a research hypothesis using much less data for validation.

## CONCLUSION

Repli-Sim is the first model of DNA replication which allows for a stochastic replication fork speed. We have shown that including stochastic replication speed is a key innovation allowing correct reconstruction of distances travelled by replication forks both in wild-type cells and in a condition when replication stress is induced. We also proposed an empirical modification to the replication fork speed, promoting completion of replication in a timely manner.

## Supporting information

Unpublished DNA fiber data

## SUPPLEMENTARY DATA

Supplementary Data (DNA tracks used in Figure 5).

## MATERIALS AND METHODS

### Deriving the formula describing 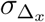

Upon origin activation, two forks are formed and elongate bidirectionally across the genome. For each specific origin, two forks replicate a distance of DNA, called DNA tracks (Δ*x*). For an origin in a cell population, during S-phase at time *t*_*exp*_ measured from G1, the distribution of Δ*x* is derived considering the following assumptions:

1. The firing time of the origin, taken as the initial time (*t*_0_), is derived from a normal distribution with a mean firing time *µ*_*t*_, specific to that origin reproducible from experimental data, and standard deviation *σ*_*t*_:

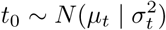
2. Individual forks have different speeds, however the speed of each fork is derived from the same probability distribution with a mean speed, *µ*_*v*_, equivalent to the average fork speed observed from experimental data, and standard deviation *σ*_*v*_:

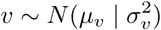

Considering the relation Δ*x* = *v* · Δ*t*, using the distribution function of Δ*t* and *v*, the distribution function of Δ*x* for each origin can be derived as follows:

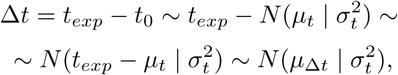

where *µ*_Δ*t*_ = *t*_*exp*_ − *µ*_*t*_.

From the other side, assuming *σ*_*t*_*σ*_*v*_ ≪ *µ*_Δ*t*_*µ*_*v*_ ([45]),

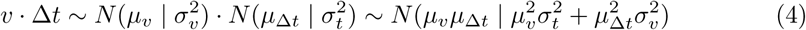

Considering Eq(4) and taking into account the assumption 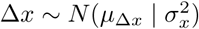, we have:

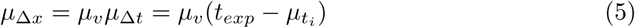

and

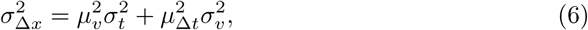

which shows that variations in DNA track distributions is dependent on variations in firing time of the origins of replication as well as variations in the fork speed progression.

### Deriving mean firing time from experimental data

For hydroxyurea-treated wild-type yeast cells, the mean firing time of individual origins is inferred from DNA copy number BrdU-labeled microarray experimental data available in [32]. At each individual origin the distribution of DNA tracks (Δ*x*) is determined and used to derive the mean firing time as follows:

1. To normalize the distribution of DNA tracks measured from BrdU experimental data, the BrdU micro-array DNA copy number of ARS305 is used and normalized to give the same efficiency as derived from its DNA copy number from quantitative PCR experiment.
2. The normalized distribution of DNA copy number is used to derive the probability distribution function for each individual origin with a p value for each DNA track length as shown in Fig8, from which *µ*_Δ*x*_ is derived.
3. Mean firing time of each origin is assumed to be individual to that origin, however variation of firing time from the mean (*σ*_*t*_) is the same for all the origins and taken as a global parameter. The firing time of *i*^*th*^ origin, (*t*_0_), is derived from the following normal distribution:

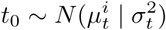
4. Individual forks have different speeds, however the speed of each fork is derived from the same probability distribution with an average speed *µ*_*v*_ and standard deviation *σ*_*v*_:

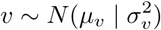

Considering the relation Δ*x* = *v* · Δ*t*, and taking into account Eq. 5, knowing the distribution function of Δ*x*, the mean firing time for each individual origin is derived as follows:

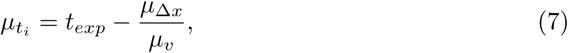

which is used in our simulations to infer the mean firing time by implementing *t*_*exp*_ and *µ*_Δ*x*_ while *µ*_*v*_ is the parameter, which is adjusted in the simulation through parameter selection in the genetic algorithm.

### Parameter selection

For parameter selection, we used a genetic algorithm to minimize the sum of the square of differences between the distribution of DNA tracks from both experimental data and simulation results. We used a population of 5000 sets of parameters, run in parallel using the open source implementation OpenMP over 32 threads. For each condition, a number of best sets of parameters was selected among which the one is chosen with a more similar total number of active origins compared to the experimental data.

### Experimental data

Throughout the analysis, three different experimental data sets ([12, 13, 23]) are utilized with the list of origins of replication detected in each individual experiment. The origins used are consistent with the OriDB database [5]. The DNA fiber data for wt cells collected during S phase, kindly provided by Philippe Pasero, are used for model selection.

## ACKNOWLEDGEMENTS

The authors are grateful to Philippe Pasero for providing DNA track data used in Fig. 5, to Philippe Pasero, John Bechhoefer, Benjamin Pardo, Yingjie Zhu, and Cheng Zhang for helpful discussions and to Heather Lander of the Sealy Center for Structural Biology and Molecular Biophysics at UTMB, for editorial services for the manuscript

